# *BRCA2* abrogation triggers innate immune responses potentiated by treatment with PARP inhibitors

**DOI:** 10.1101/490268

**Authors:** Timo Reisländer, Emilia Puig Lombardi, Florian J. Groelly, Ana Miar, Manuela Porru, Benjamin Wright, Helen Lockstone, Annamaria Biroccio, Adrian Harris, Arturo Londono, Madalena Tarsounas

**Author notes:** These authors contributed equally to this study.

## Abstract

Heterozygous germline mutations in *BRCA2* predispose to breast and ovarian cancer. Contrary to non-cancerous cells, where *BRCA2* deletion causes cell cycle arrest or cell death, *BRCA2* inactivation in tumors is associated with uncontrolled cell proliferation. We set out to investigate this conundrum by exploring modalities of cell adaptation to loss of BRCA2 and focused on genome-wide transcriptome alterations. Human cells in which BRCA2 expression was inhibited using a doxycycline (DOX)-inducible shRNA for 4 or 28 days were subjected to RNA-seq analyses. Gene sets differentially expressed in BRCA2-deficient versus -proficient cells revealed a biphasic response to *BRCA2* abrogation. The early, acute response consisted of downregulation of genes involved in cell cycle progression, DNA replication and repair and was associated with cell cycle arrest in G1. Surprisingly, the late, chronic response consisted predominantly of upregulation of innate immune response genes controlled by interferon. Activation of the cGAS-STING pathway detected in these cells further substantiated the concept that long-term *BRCA2* abrogation triggers cell-intrinsic immune signaling. Importantly, we found that treatment with PARP inhibitors stimulated the interferon response in cells and tumors lacking BRCA2. We propose that PARP inhibitors may suppress growth of BRCA2-deficient cells and tumors, in part, by activating interferon signaling.

---

BRCA2 tumor suppressor plays key roles in cell physiology by promoting DNA replication and DNA double-stranded breaks (DSBs) repair via homologous recombination^1^. The latter is well-characterized biochemically and relies on BRCA2 loading the RAD51 recombinase at sites of DSBs that have been processed by resection. Assembly of the RAD51 nucleoprotein filament facilitates the search and subsequent invasion of homologous DNA, which acts as a template for the repair reaction^2^. Whilst BRCA2 function in DNA repair has been studied in the context of DNA damage inflicted by exposure to genotoxic agents (e.g. ionizing radiation, DNA cross linking agents), the role of BRCA2 in replication is intrinsic to cell physiology. In the absence of external challenges, loss of BRCA2 triggers a significant decrease in replication fork progression and a high frequency of stalled forks^3,4^. A subset of these forks collapse and are converted to DSBs which, due to compromised HR repair, accumulate in BRCA2-deficient cells. Moreover, nucleolytic degradation of the stalled forks can occur upon extensive replicative arrest^5,6^. Conceivably, this acute perturbation of DNA replication triggered by inactivation of BRCA2 could lead to rampant genomic instability, which is lethal or severely obstructs cell proliferation in primary human cells^7^. Likewise, disruption of *Brca2* gene in mice is embryonically lethal^8–10^. Mechanisms of replication stress and DNA damage tolerance mediate cellular adaptation to chronic loss of BRCA2, enable cells to survive and ultimately underlie their tumorigenic potential. Consistent with this, loss of *BRCA2* occurs in tumors and is thought to promote tumorigenesis, whilst *BRCA2* heterozygous germline mutations increase susceptibility to breast and ovarian cancer, as well as other cancers^11–13^.

Here we investigate the possibility that transcriptional alterations provide modalities of cell adaptation to loss of BRCA2, thus preventing cell death or proliferative arrest. We characterized the transcriptome of BRCA2-deficient cells, using a doxycycline (DOX)-inducible shRNA to inhibit BRCA2 expression in human non-small cell lung carcinoma H1299 cells and invasive ductal breast cancer MDA-MB-231 cells. RNA-sequencing (RNA-seq) analyses conducted after 4 and 28 days of DOX-induced BRCA2 depletion enabled us to monitor the dynamics of gene expression and to identify substantial transcriptional alterations from early to late stages of BRCA2 inactivation. In the short term, we observed downregulation of cell cycle DNA replication and repair genes, which correlated with marked accumulation of BRCA2-deficient cells in G1. In the long-term, we found that cell cycle re-entry occurred concomitantly with upregulation of a set of genes involved in the innate immune response and controlled by interferon signaling^14^. A similar set of genes was upregulated in *BRCA2*-deleted primary ovarian tumors. These results support the concept that inducible BRCA2 inactivation in cultured cells recapitulates cellular changes associated with loss of BRCA2 during tumorigenesis and could provide clues to mechanisms of cellular adaptation in tumors or tumor vulnerabilities. Importantly, treatment with PARP inhibitors (olaparib, talazoparib) stimulated upregulation of the interferon signaling genes, which may contribute, in part, to the toxicity of PARP inhibitors to BRCA2-deficient cells and tumors.

## RESULTS

### BRCA2 inactivation elicits changes in gene expression

Previous studies have implicated BRCA2 in gene transcription via interaction with EMSY^15^ and have reported differential gene expression between *BRCA2*-deleted and wild type cells using expression microarrays^16^. *BRCA2*-deleted cells are terminally adapted and this precluded analysis of progressive changes associated BRCA2 inactivation. Here we used RNA-seq analyses of gene expression to determine how the transcriptome of BRCA2-deficient cells is modulated over time. We cultured H1299 and MDA-MB-231 human cells carrying a DOX-inducible BRCA2 shRNA cassette in the presence or absence of 2 μg/mL DOX for 28 days and collected samples for RNA-seq analysis after 4 and 28 days of treatment (Fig. 1a). BRCA2 expression was effectively suppressed by DOX exposure in the short-term (4 days), as well as in the long-term (28 days), as indicated by immunoblotting of cell extracts prepared at these time points (Fig. 1b).

**Figure 1.**
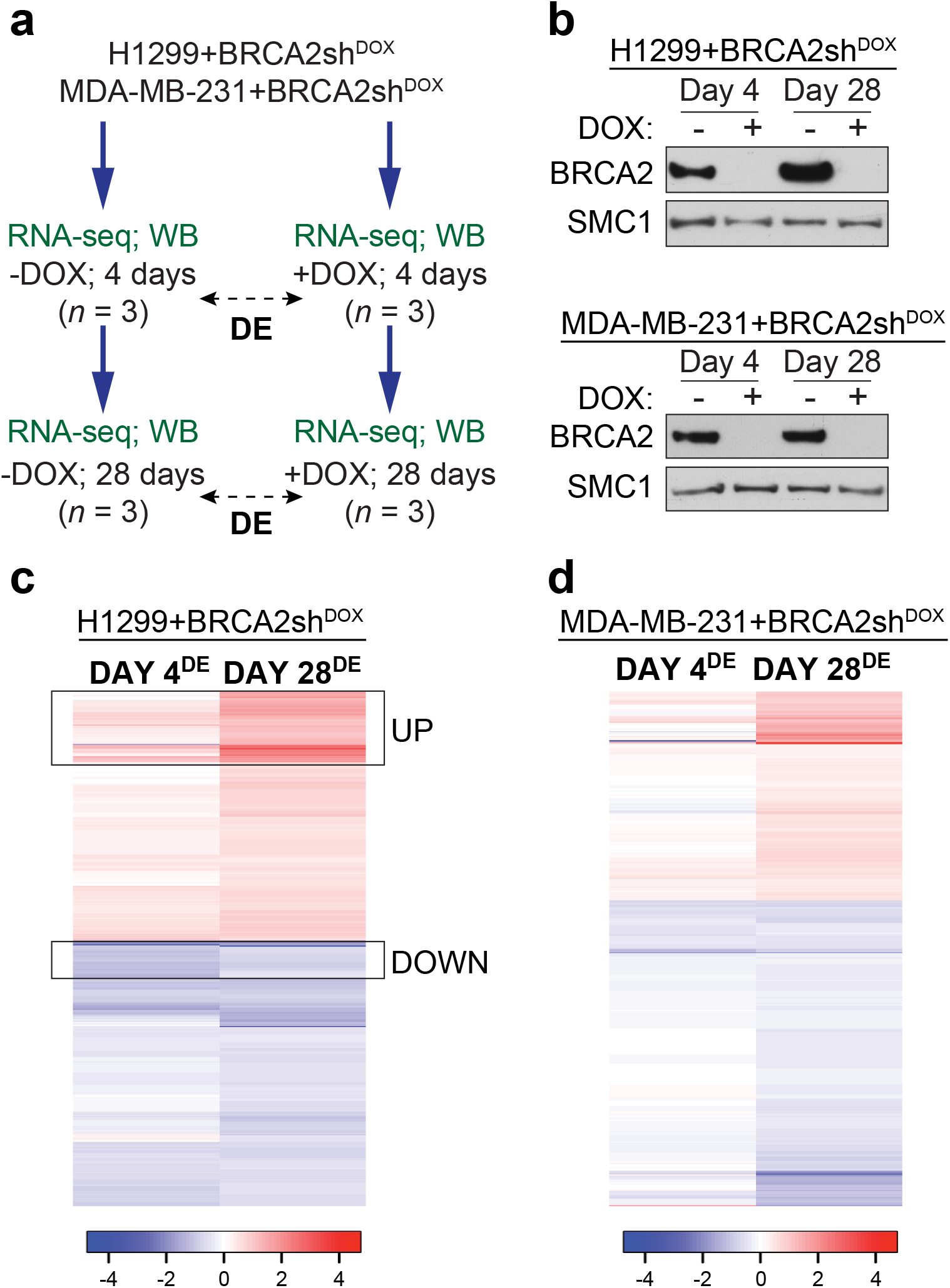
Transcriptome changes during the transition from short-term, acute to long-term, chronic *BRCA2* inactivation in human cells. (**a**) Human H1299 and MDA-MB-231 cells carrying a doxycycline (DOX)-inducible BRCA2 shRNA were grown in the presence or absence of 2 μg/mL DOX for 4 or 28 days before processing for RNA-seq and Western blot analyses. (**b**) Whole-cell extracts prepared after 4 or 28 days of DOX treatment were immunoblotted as indicated. SMC1 was used as a loading control. (**c, d**) RNA-seq analyses of cells treated as in (**a**) identify transcriptional alterations specific to BRCA2-deficiency (FDR < 0.05) after 4 and 28 days of DOX treatment. Heatmaps depict Log2(Fold Change) of top 480 genes differentially expressed in +DOX versus −DOX cells, after 4 or 28 days of DOX treatment. Boxes indicate a subset of genes downregulated at day 4 (DOWN) or upregulated at day 28 (UP) in BRCA2-deficient H1299 cells. *n* = 3 independent experiments; DE, differential expression.

Before proceeding with RNA-seq analyses of BRCA2-deficient cells, we performed control experiments in which we addressed whether DOX alone could induce transcriptional changes under our experimental conditions. Parental H1299 and MDA-MB-231 cells were incubated with 2 μg/mL DOX for 4 and 28 days, before collecting samples for RNA-seq analyses (Supplementary Fig. 1a,b). Hierarchical clustering of sample-to-sample Euclidean distances based on the RNA-seq data showed no clear clustering of samples treated with DOX (*n* = 3 independent experiments) or untreated samples (*n* = 3 independent experiments) in either cell line, indicative of insignificant alterations between the −DOX and +DOX samples. Moreover, differential gene expression analyses with false discovery rate (FDR) of 5% revealed that only 2 genes were differentially expressed in DOX-treated versus untreated H1299 cells at day 4 and none at day 28 of DOX treatment (Supplementary Fig. 1c). The corresponding numbers of deregulated genes in MDA-MB-231 were 3 and 7, respectively (Supplementary Fig. 1d). These results demonstrate that DOX treatment for 4 or 28 days has a negligible impact on gene expression of parental cells lacking BRCA2 shRNA.

In contrast to this, DOX treatment inflicted substantial changes on the transcriptome of H1299 and MDA-MB-231 cells carrying a DOX-inducible BRCA2 shRNA. Hierarchical clustering of sample-to-sample Euclidean distances showed a clear distinction between +DOX (*n* = 3 independent experiments) and −DOX samples (*n* = 3 independent experiments) in both cell lines (Supplementary Fig. 2a,b), indicative of specific, substantial differences in the transcriptome upon BRCA2 abrogation. Consistent with this, DOX-inducible BRCA2 depletion in H1299 cells triggered significant alterations (FDR < 0.05, no fold-change filter) in the expression of 1,363 genes at day 4 and of 479 genes at day 28 of DOX treatment (Supplementary Fig. 2c and Supplementary Table 1). The corresponding numbers of deregulated genes in MDA-MB-231 cells were 187 and 480, respectively (Supplementary Fig. 2d and Supplementary Table 2), suggesting that BRCA2 abrogation had a less pronounced effect on gene expression in the short-term (4 days) in this cell lines.

Next we conducted supervised clustering analyses of the top 480 genes differentially expressed (FDR < 0.05, no fold-change filter) in H1299 and MDA-MB-231 cells upon DOX-induced BRCA2 inactivation (Fig. 1c,d). We observed a clear variation in the differential transcription profiles between day 4 and day 28 of DOX treatment in both cell lines, which suggests that the transcriptome of BRCA2-deficient cells undergoes dynamic changes with time. The heatmaps showing the expression profiles of H1299 cells at replicate level are shown in Supplementary Fig. 3.

### Distinct pathways are deregulated in the short-term versus long-term BRCA2 inactivation

We proceeded to identify which gene sets are differentially expressed in BRCA2-deficient versus -proficient cells, at 4 and 28 days after shRNA induction. We focused on H1299 cells because DOX treatment had a high impact on gene expression in this cell line. We used stringent conditions (FDR < 0.05; Log_2_Fold Change > |0.5|) which enabled us to identify deregulated genes with high confidence (Fig. 2a,b). At day 4 of DOX treatment, this analysis identified 574 genes (42% of deregulated genes) significantly downregulated (Log2Fold Change < -0.5; Supplementary Table 1) and 147 genes (11% of deregulated genes) significantly upregulated (Log2Fold Change > 0.5) in BRCA2-deficient cells. At day 28, there were 194 genes (40%) significantly downregulated (Log2Fold Change < -0.5; Supplementary Table 2) and 213 genes (44%) significantly upregulated (Log2Fold Change > 0.5;) in BRCA2-deficient H1299 cells.

**Figure 2.**
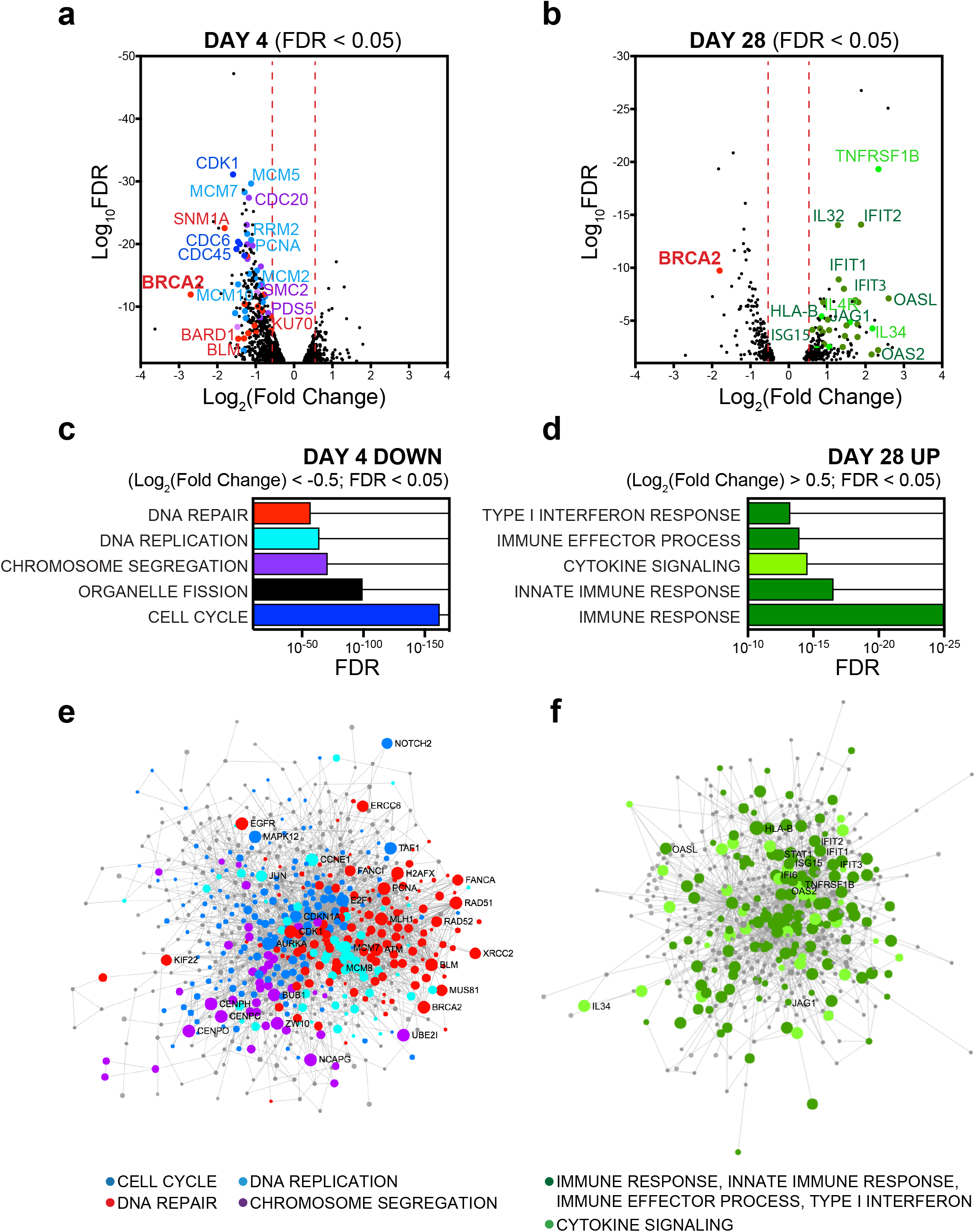
Pathway deregulation in BRCA2-deficient H1299 cells during short-term (4 days) or long-term (28 days) *BRCA2* inactivation using a DOX-inducible shRNA. (**a, b**) Volcano plot of genes differentially expressed (FDR < 0.05) in BRCA2-deficient versus BRCA2-proficient H1299 cells, after DOX treatment for 4 (a) or 28 days (**b**). A subset of the genes significantly downregulated (Log2(Fold Change) < -0.5) after 4 days or significantly upregulated (Log_2_(Fold Change) > 0.5) after 28 days of DOX treatment is shown. (**c, d**) Gene set enrichment analysis based on functional annotation (Gene Ontology - Biological Process database) of genes downregulated after 4 days (**c**) or upregulated after 28 days of DOX treatment (**d**). (**e, f**) First-order protein-protein interaction networks between genes downregulated after 4 days (**e**) or upregulated after 28 days of DOX treatment (**f**). Node size is proportional to the number of its connections with other nodes. *n* = 574 downregulated genes (day 4) and *n* = 213 upregulated genes (day 28) were used for network analysis, resulting in 669 and 446 nodes, respectively.

Gene set enrichment analysis of differentially expressed genes based on functional annotation (Gene Ontology - Biological Process database) showed enrichment in specific pathways (Fig. 2c,d). Genes downregulated in BRCA2-deficient cells at day 4 were mainly implicated in cell cycle, chromosome segregation, DNA repair and DNA replication and defined an early, acute response to BRCA2 inactivation. The genes upregulated at day 28 primarily mediated cytokines and immune responses. Interestingly, induction of these genes did not correlate with changes in BRCA2-deficient cell proliferation from early to late time points (data not shown).

To complement pathway mapping, we performed network analysis which showed how the 574 downregulated genes interact with each other and clustered into different pathways (Fig. 2e). We retrieved high-confidence, experimentally validated protein-protein interactions through the NetworkAnalyst platform^17^. The network was generated by mapping the significant genes to the STRING interactome^18^ and applying a search algorithm to identify first-order neighbours (proteins that directly interact with a given protein) for each of the mapped genes. We generated a highly-connected first-order network consisting of 669 nodes and 2,970 edges, with larger nodes indicating higher connectivity. For example, prominent DNA repair nodes identified key DNA repair factors including RAD51, RAD52, BML, MUS81, FANCA, MLH1, whilst main DNA replication nodes were identified by factors such as PCNA, MCM2, RRM2. This indicates concerted action of the genes downregulated at day 4 in key cellular processes, such as control of cell cycle progression and genome integrity. Similar mapping of the 213 genes upregulated at day 28 generated a network with fewer interconnected nodes (446 nodes) and 1,190 edges (Fig. 2f). Pathway analysis after addition of direct interactors amplified enrichment in immune system related pathways (hypergeometric test *p*-value < 1e^−21^). The top 5 impacted pathways were: innate immune response (77 hits, *p*-value = 3.93e^−27^), immune response (116 hits, *p*-value = 1.83e^−25^), interaction with host (61 hits, *p*-value = 1.86e^−25^), cytokine-mediated signaling (54 hits, *p*-value = 9.34e^−23^) and defense response (112 hits, *p*-value = 2.9e^−21^).

Differential gene expression analyses were also conducted in MDA-MB-231 cells carrying a DOX-inducible BRCA2 shRNA (Supplementary Fig. 4a). These revealed only 19 genes downregulated (Log2Fold Change < -0.5; Supplementary Table 1) at day 4 and 91 genes upregulated (Log2Fold Change > 0.5; Supplementary Table 2) at day 28 of DOX treatment. REACTOME biological pathway analyses showed enrichment in processes including α,β-interferon signalling and cytokine signalling in the immune system (Supplementary Fig. 4b) and were therefore analogous to the processes upregulated in H1299 cells after 28 days of DOX treatment. Moreover, we performed RNA-seq analyses of differential gene expression in *BRCA2^−/−^* versus *BRCA2^+/+^* DLD1 cells (Supplementary Fig. 5). Pathway deregulation score analyses^19^ of the REACTOME database using the top upregulated genes in *BRCA2^−/−^* DLD1 cells identified interferon signaling, cytokine signaling and immune response as the highest scoring pathways.

### Induction of innate immune response genes in BRCA2-deficient human cells

Long-term BRCA2 inactivation in H1299 and MDA-MB-231 human cells led to upregulation of common immune processes. To facilitate experimental validation for these findings, we identified common genes induced in both cell lines after 28 days of DOX treatment (Fig. 3a). We intersected the list of genes significantly upregulated in H1299 and MDA-MB-231 cells at day 28 (FDR < 0.05) and further filtered the high-confidence hits using Log2Fold Change > 0.3. This analysis identified 28 genes upregulated in both cell lines (Table 1), which showed enrichment in Gene Ontology processes including cytokine signaling, type I interferon response and immune response (Fig. 3b).

**Figure 3.**
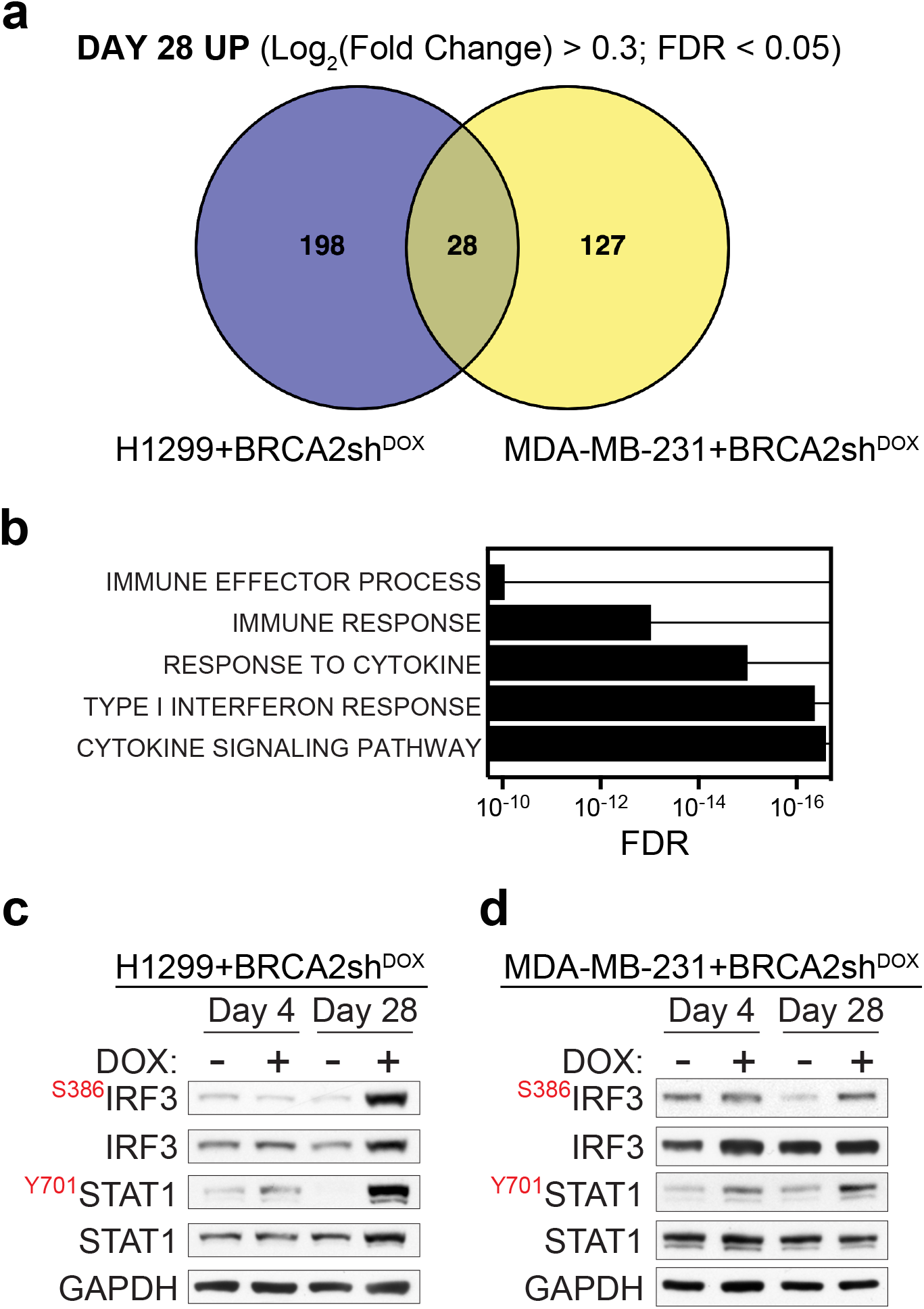
Upregulation of innate immune response genes in BRCA2-deficient human cells. (**a**) Venn diagram of common genes upregulated in H1299 and MDA-MB-231 human cells, upon chronic (28 days) DOX-induced BRCA2 depletion. (**b**) Gene set enrichment analysis (Gene Ontology - Biological Process database) of the 28 genes upregulated upon chronic BRCA2 inactivation in both H1299 and MDA-MB-231 cells. (**c, d**) Whole-cell extracts prepared after 4 and 28 days of DOX treatment were immunoblotted as indicated. GAPDH was used as a loading control. Phosphorylation site is indicated in red.

In BRCA2-deficient cells, replication fork instability and nucleolytic degradation inflict replication-associated DNA damage, which conceivably cause cytosolic DNA accumulation and activate innate immune responses^14^. Phosphorylation of STAT1 at Tyr701 is routinely used as a marker for activation of cytokine signaling^20^ including interferon type I^21^ in response to cytosolic DNA. Consistent with this, we observed enhanced STAT1 Tyr701 phosphorylation (Fig. 3c) in BRCA2-deficient relative to BRCA2-proficient H1299 cells, after 28 days of DOX treatment (Fig. 3c). IRF3 phosphorylation at Ser386, indicative of its nuclear translocation^22^, was also induced by long-term BRCA2 depletion. A similar response was triggered in MDA-MB-231 by BRCA2 depletion for 28 days (Fig. 3d). Thus, BRCA2 inhibition in human cells activates DNA damage responses and stimulates innate immune gene expression, as previously reported for DNA damaging agents^20,21^ and SAMHD1 inactivation^23^.

### Validation of innate immune response activation upon chronic *BRCA2* inhibition

Induction of interferon response gene expression identified using RNA-seq analyses was validated using quantitative RT-PCR (Fig. 4a). mRNA levels of interferon-stimulated genes (*IFIT1, IFIT2, IFIT3, IFI6, OAS1, OAS2, ISG15*) and cytokine signaling gene (*TNFRSF1B*) were significantly upregulated upon BRCA2 inactivation for 28 days, whilst no significant changes were detected after 4 days of DOX treatment. Consistent with the concept that interferon signaling triggers JAK/STAT-dependent gene expression, we found that STAT1 siRNA-mediated depletion decreased the mRNA levels of interferon-stimulated genes (Supplementary Fig. 6a). Moreover, we found that a subset of these genes were also upregulated in *BRCA2*-deleted ovarian tumors (Fig. 4b). The Cancer Genome Atlas (TCGA) ovarian serous cystadenocarcinoma mRNA expression data for 309 cases were used to classify tumors relative to their *BRCA2* mRNA levels. The 3rd highest quantile of mRNA expression defined the group of *BRCA2*^high^ tumors (*n* = 71). The cases carrying *BRCA2* deep deletions (*n* = 4) were classified as *BRCA2*^del^. Out of the list of 8 genes validated by quantitative RT-PCR, the 7 interferon-stimulated genes *(IFIT1, IFIT2, IFIT3, IFI6, OAS1, OAS2, ISG15)* had higher mRNA levels in *BRCA2*^del^ compared to *BRCA2*^high^ tumors. These results suggested that upregulation of this subset of innate immune response genes upon chronic inactivation of *BRCA2* observed *in vitro* is recapitulated in ovarian human tumors. Notably, four of the interferon-stimulated genes (*IFIT1, IFIT2, OAS2, ISG15*) were also upregulated in the MDA-MB-231 cells lacking BRCA2 after 28 days of DOX treatment (Supplementary Fig. 7a), albeit at lower levels than in H1299 cells. This result may reflect transcriptional regulation specific to the breast tumor from which this cell line originates. Supporting this observation, TCGA breast invasive carcinoma tumor data analysis revealed insignificant differences in the expression of the same gene set between BRCA2-deleted tumors and those with high *BRCA2* mRNA expression (data not shown).

**Figure 4.**
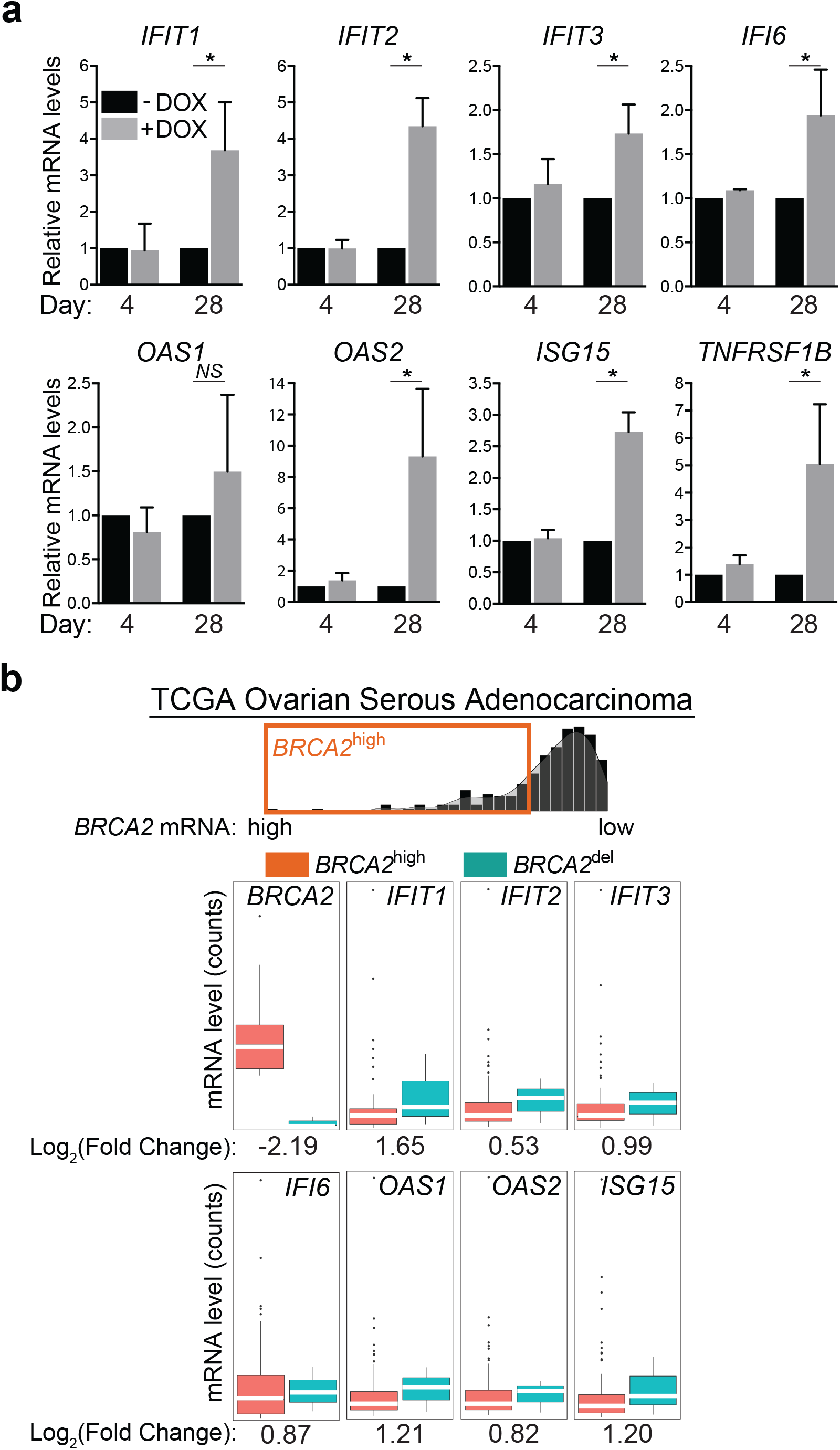
Induction of interferon-stimulated genes in BRCA2-deficient cells and tumors. (**a**) mRNA levels of indicated genes were determined using quantitative RT-PCR analyses and were expressed relative to the housekeeping gene *RPL11* and to untreated (-DOX) control cells (2^−ΔΔCT^). Error bars represent SD of *n* = 3 independent experiments, each performed in triplicate. *NS,p* > 0.05; *,*p* < 0.05 (paired two-tailed *t* test). (**b**) Upregulation of innate immune response genes in BRCA2-deleted ovarian tumors (*n* = 4) versus tumors with high *BRCA2* mRNA expression (*n* = 71). Dots in graphs represent individual tumors. Middle line (white), median; box limits 25 and 75 percentiles; whiskers, minimum and maximum values.

Next we evaluated the dynamics of immune response gene expression by monitoring changes in mRNA levels with time (Fig. 5a). Cells carrying a DOX-inducible shRNA against BRCA2 were treated with DOX for 28 days. Gene expression was measured at 2-day intervals using quantitative RT-PCR. mRNA levels for all genes monitored in this manner increased gradually during chronic BRCA2 inactivation in H1299 cells. Immune response gene upregulation was also detected over time in MDA-MB-231 cells (Supplementary Fig. 7b), albeit to lower levels than in H1299. Additionally, DOX-induced BRCA2 depletion triggered STAT1 phosphorylation at Tyr701 and IRF3 phosphorylation at Ser386 (Fig. 5b). These preceded upregulation of the innate immune response genes, consistent with IRF3 and STAT1 promoting transcription of the interferon-stimulated genes. Interestingly, STING protein levels were also markedly increased upon BRCA2 abrogation, while no significant change was detected in its mRNA levels. This may reflect stabilization of this protein, possibly by inhibition of its proteolytic degradation^24^.

**Figure 5.**
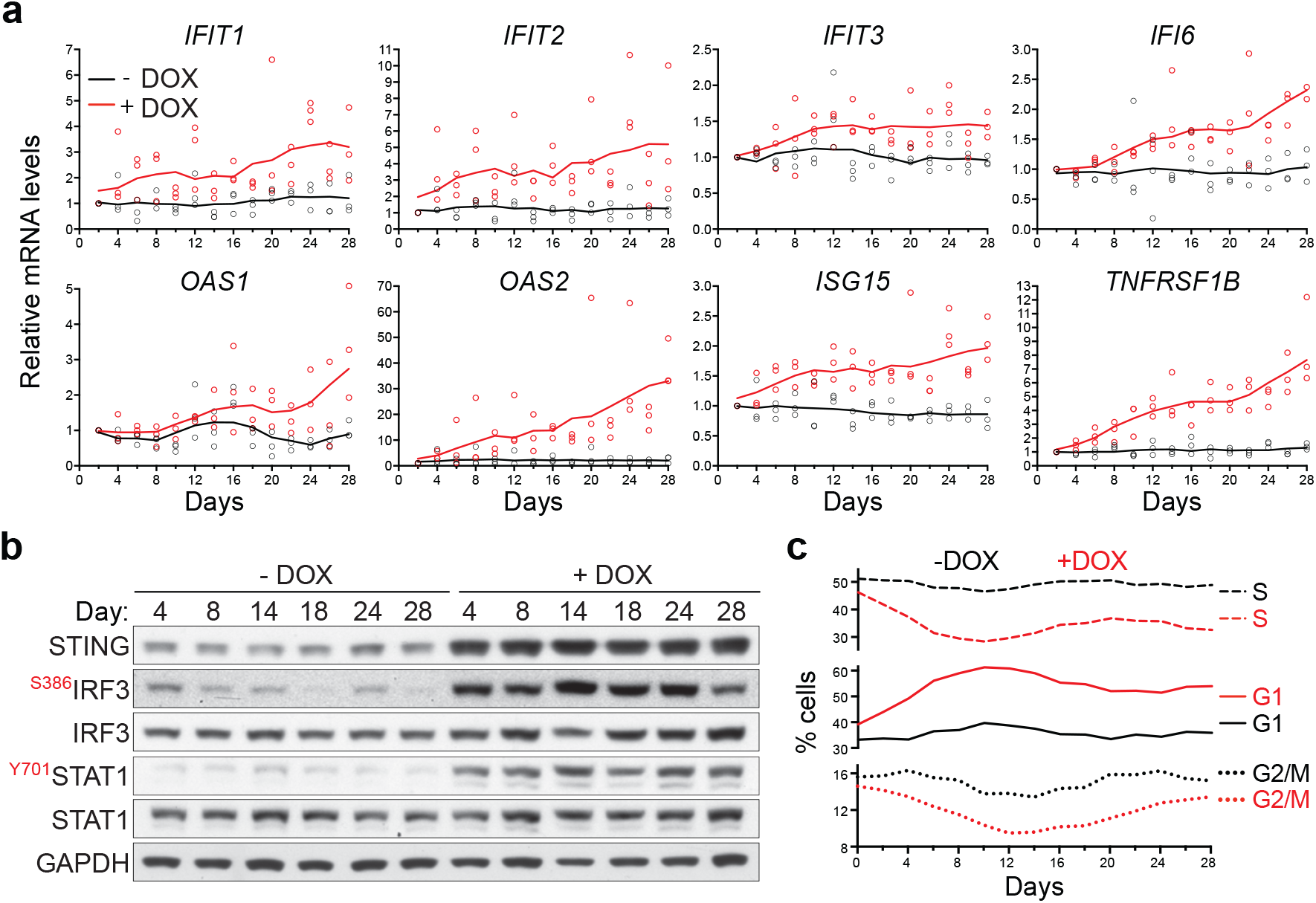
Time course of innate immune response activation in BRCA2-deficient cells. (**a**) H1299 cells expressing a DOX-inducible BRCA2 shRNA were grown in the presence or absence of DOX for 28 days. Cells collected every two days were subjected to quantitative RT-PCR analyses using primers specific for the indicated genes. mRNA levels were expressed relative to the gene encoding β-actin and to untreated (-DOX) control cells at every time point. *n* = 3 independent experiments, each in performed in triplicate. Each dot represents a single replicate. (**b**) Whole-cell extracts prepared at the indicated times after DOX addition were immunoblotted as shown. GAPDH was used as a loading control. Phosphorylation sites are indicated in red. (**c**) Cells treated as in (**a**) were pulse-labelled with EdU for 30 minutes. Frequency of cells in G1, S and G2/M stages of the cell cycle were determined using FACS analyses of EdU-labeled cells.

In the course of these experiments we also addressed the possibility that loss of BRCA2 impacts on cell cycle progression. FACS analyses of EdU-pulsed H1299 cells revealed pronounced alterations in the cell cycle profile of BRCA2-deficient cells during the transition from acute to chronic responses (Fig. 5c). Cells accumulated in G1 in response to acute BRCA2 inactivation. Concomitantly, we detected decreased levels of cells in S and G2/M, indicative of cell cycle arrest in G1. This may be a consequence of transcriptional downregulation of genes controlling cell cycle and DNA replication demonstrated by RNA-seq analyses of BRCA2-deficient cells (Fig. 2a,c). The frequency of G1 cells reached a peak at day 12 and subsequently diminished, without however reaching the level of BRCA2-proficient cells. The percentage of S-phase cells remained lower in cells lacking BRCA2 relative to wild type for the course of the experiment. These results are consistent with the observation that BRCA2-deficient cells proliferate more slowly than BRCA2-proficient cells (data not shown).

### PARP inhibitors potentiate the expression of innate immune response genes in BRCA-deficient human cells and tumors

Small molecule inhibitors of poly [ADP-ribose] polymerase (PARP) have been recently approved for clinical treatment of BRCA1- and BRCA2-deficient ovarian and breast cancers^25,26^. The molecular mechanisms of selective targeting of BRCA-deficient cells and tumors by PARP inhibitors has been investigated extensively. Trapping of the PARP1 on DNA ends obstructs replication fork progression leading to accumulation of single strand breaks, which are converted into lethal DSB lesions^27–30^.

Induction of DNA damage by PARP inhibitors prompted us to investigate whether treatment of BRCA2-deficient cells with these drugs could also impact on the interferon response. To address this, we treated H1299 cells with DOX for 4 days to induce BRCA2 depletion. Under these conditions, we did not detect upregulation of interferon-stimulated genes using quantitative RT-PCR (Fig. 4a, Fig. 5a). However, incubation with 1 or 10 μM olaparib for 24 hours induced a dose-dependent increase in the mRNA levels of interferon-stimulated genes (Fig. 6a). We concluded that olaparib induced innate immune responses even after 4 days of BRCA2 depletion.

**Figure 6.**
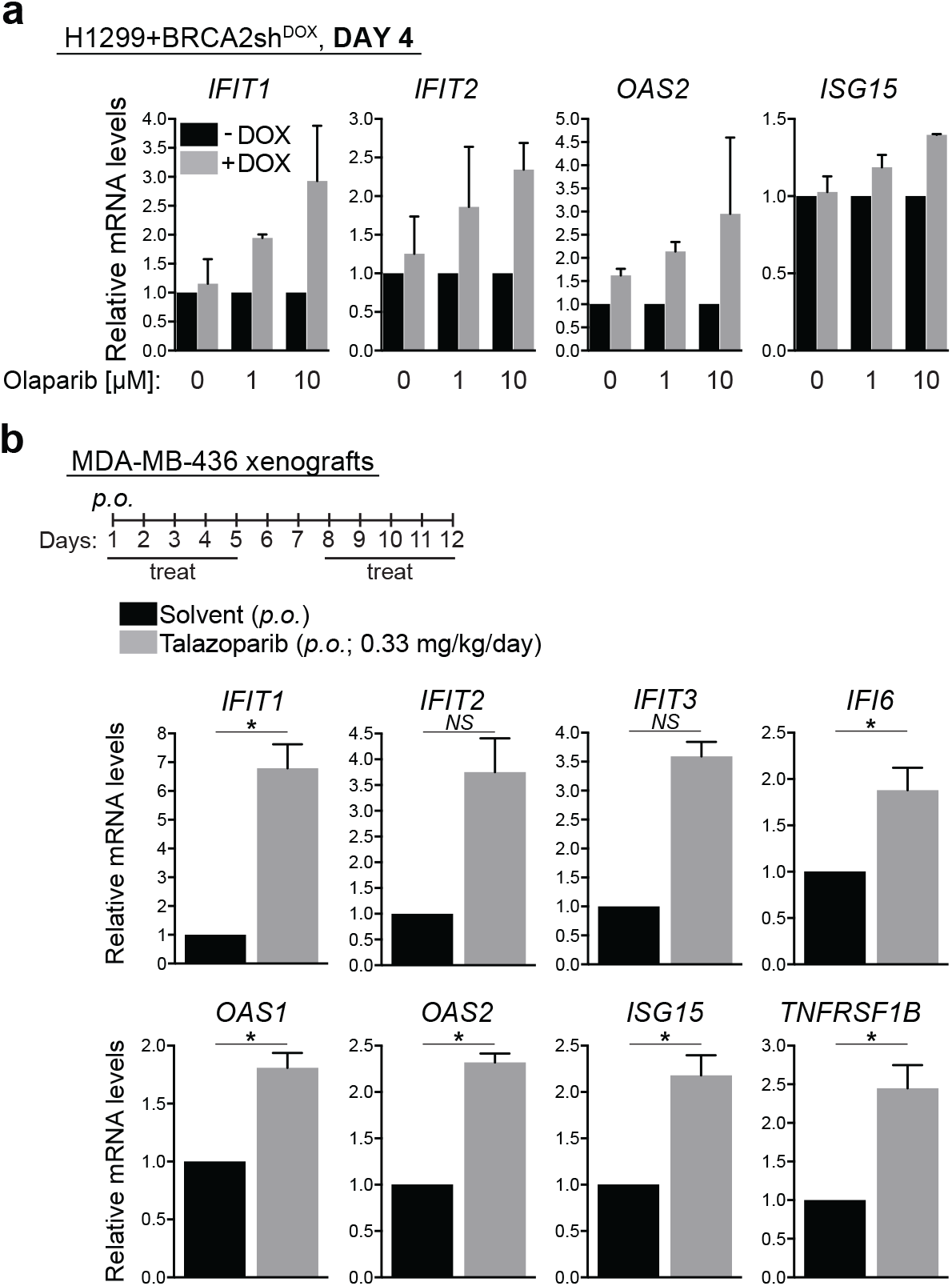
Effect of PARP inhibitors on the interferon-stimulated gene expression triggered by BRCA2 inactivation. (**a**) H1299 cells expressing a DOX-inducible BRCA2 shRNA were grown in the presence or absence of DOX for 4 days. Olaparib (1 or 10 μM) was added for the last 24 hours of DOX treatment, followed by quantitative RT-PCR analyses using primers specific for the indicated genes. mRNA levels were expressed relative to the gene encoding β-actin and to untreated (-DOX) control cells (2^−ΔΔCT^). (**b**) BRCA1-deficient MDA-MB-436 human cells (1 x 10^6^) were injected into the mammary fat pad of CB17/SCID female mice. Tumor-bearing mice were treated on the indicated days with PARP inhibitor talazoparib or with solvent control, both orally administered (*p.o*.). Total RNA was prepared from tumors collected at the end of the treatment. mRNA levels of indicated genes were determined using quantitative RT-PCR analyses and were expressed relative to the gene encoding β-actin and to solvent-treated control tumors (2^−ΔΔCT^). Error bars represent SD of *n* = 3 tumors, for which quantitative RT-PCR reaction was performed in triplicate. *NS, p* > 0.05; *, *p* < 0.05 (unpaired two-tailed *t* test).

Next, we investigated whether the selective toxicity of olaparib against BRCA2-deficient cells could be mediated by the interferon response. Interferon signaling is attenuated when STAT1 expression is inhibited using siRNA (Supplementary Fig. 6a). Thus, we treated BRCA2-proficient and -deficient H1299 cells with an siRNA against STAT1, before exposing them to 10 μM olaparib for 72 hours. We observed that cells in which BRCA2 was abrogated for 4 days were sensitive to olaparib and that STAT1 inactivation rescued their sensitivity (Supplementary Fig. 6b). This indicates that the selective elimination of BRCA2-deficient cells by olaparib may be in part mediated by induction of innate immune responses. In contrast, STAT1 inhibition in BRCA2-proficient cells increased susceptibility to olaparib. Moreover, olaparib sensitivity of cells in which BRCA2 was chronically inactivated (28 days) were not affected by loss of STAT1, consistent with a role for STAT1 during the early stages of BRCA2 inactivation (Supplementary Fig. 6c).

To address whether the innate immune response triggered by BRCA inactivation *in vitro* can be recapitulated *in vivo*, in the tumor context, we established orthotopic xenograft tumors by injecting BRCA-compromised MDA-MB-436 mammary breast cancer cells^31^ into the mammary fat pad of CB17/SCID mice (Fig. 6b). Tumor treatment with the PARP inhibitor talazoparib triggered interferon-stimulated gene expression, as demonstrated by quantitative RT-PCR analysis of tumor RNA. Thus, PARP inhibitors elicit innate immune responses in BRCA-deficient cells and tumors.

## DISCUSSION

Although not lethal, BRCA2 inactivation in cancer-derived human cells significantly slows down cell proliferation. This indicates that cells can adapt to loss of BRCA2 and to the perturbations in cell physiology this entails. Here we explored the changes in transcriptome associated with BRCA2 abrogation and defined distinct transcriptional alterations, evolving from an early, acute response, to a late, chronic response. The former was characterized primarily by decreased mRNA levels of genes required for cell cycle progression, chromosome segregation, DNA repair and replication, consistent with previously reported roles for BRCA2 in replication fork stability and DNA repair via homologous recombination reactions^1^. EdU incorporation analyses revealed that these transcriptional changes are associated with accumulation of BRCA2-deficient cells in G1 and with low levels of entry/progression through S and G2/M phases of the cell cycle. Slowing down these processes may be conducive for activation of alternative, BRCA2-independent mechanisms of fork protection and re-start, which, in the long-term, sustain cell survival.

The late, chronic response to BRCA2 inactivation consisted of upregulation of interferon-regulated innate immune response genes. best. Initially identified as a mechanism of defense against pathogens by blocking viral infections and priming adaptive immune responses^32^, the innate immune response can also be enlisted to counteract tumor progression^14^. The response operates on the premise that free genomic DNA evicted into the cytoplasm as a consequence of the genomic instability intrinsic to most tumors is recognized as pathogenic by cGAS and triggers STING activation (Fig. 7). This, in turn, facilitates phosphorylation and nuclear translocation of interferon response factors (e.g. IRF3), which drive transcription of the interferon response genes. Secreted interferon acts similarly to other cytokines to activate signaling pathways, in particular the JAK/STAT1 pathway, within the same cell (autocrine response) or in adjacent cells (paracrine response), resulting in induction of interferon-stimulated genes. Expression of these genes can also be triggered by ionizing radiation^20^ and chemotherapeutic drugs (e.g. cisplatin, MMC)^21^ and requires passage through mitosis with associated micronuclei formation^20,33^.

**Figure 7.**
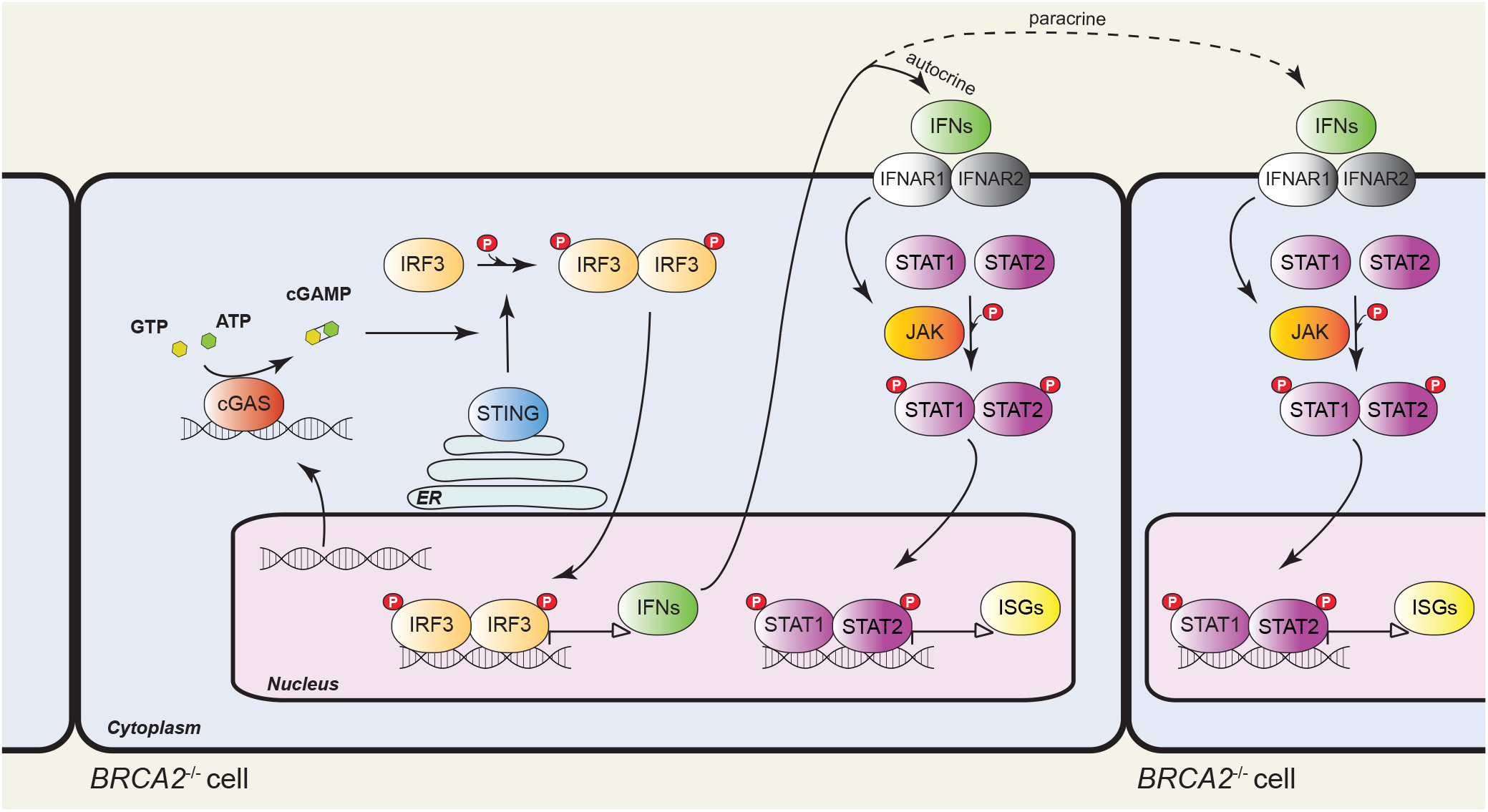
Model of innate immune response activation in BRCA2-deficient cells. Intrinsic replication stress and unrepaired DSBs cause genomic instability and DNA translocation to the cytosol, where it activates cyclic GMP-AMP synthase (cGAS), a cytosolic DNA sensor. cGAS catalyses the synthesis of cyclic GMP-AMP (cGAMP), which binds to and activates STING (stimulator of interferon genes). STING promotes TBK1-dependent phosphorylation of interferon response factor 3 (IRF3), which translocates into the nucleus as a homodimer to induce transcription of type I interferon (IFN). Secreted IFN binds to transmembrane receptors and elicits autocrine and paracrine signaling, both mediated by the JAK-STAT pathway. Janus kinases (JAKs) phosphorylate signal transducer and activator of transcription 1 (STAT1) and 2 (STAT2), which enter the nucleus as a heterodimer to initiate transcription of IFN-stimulated genes (ISGs). ER, endoplasmic reticulum.

Our results demonstrate that the mRNA levels of interferon-stimulated genes are also enhanced by chronic BRCA2 inactivation. We previously reported that severe replication stress^5^ and persistent DNA damage associated with loss of BRCA2 result in aberrant chromosome segregation^4^. Conceivably, this could lead to cytosolic DNA accumulation and interferon signaling, as suggested by the phosphorylation of STAT1 and IRF3, as well as upregulation of interferon response genes observed upon chronic BRCA2 inactivation (Fig. 5a). Consistent with STAT1-dependent transcriptional activation of these genes (Supplementary Fig. 6a), our results demonstrate that STAT1 Tyr701 phosphorylation, required for its nuclear translocation and activation, is detectable from the early stages of BRCA2 depletion (Fig. 5b).

How the innate immune response activation impacts on the long-term survival of BRCA2-deficient cells and tumors remains to be elucidated. In our *in vitro* system, the interferon response induced upon chronic BRCA2 inactivation is associated with partial restoration of cell cycle progression (Fig. 5c). Upregulation of immune signaling factors known to promote cell proliferation (e.g. TNFRSF1B, also known as TNFR2 or p75 ^34^) could account for the ability of BRCA2-deficient cells to survive. However, prolonged cell survival in the absence of BRCA2 may potentiate the innate immune response, as successive rounds of DNA replication and chromosome segregation are likely to augment their genomic instability. In the tumor context, ensuing interferon responses may stimulate cytotoxic T-cell activation, analogous to the STING-dependent defense mechanisms elicited by viral infection^35^, which could be onco-suppressive. Studies using mouse models of tumorigenesis driven by *Brca2* abrogation^36^ must be conducted in order to evaluate this possibility. Conversely, it will be interesting to determine whether BRCA2-deficient tumors that became resistant to chemotherapy have also acquired the ability to escape immune detection.

A recent study has shown that PARP inhibitors inflict DNA damage and promote cGAS/STING pathway activation *in vitro*, independently of BRCA status^37^. Our results establish that olaparib accelerates upregulation of innate immune response genes specifically in BRCA-deficient cells and tumors. Conceivably, the intrinsically high levels of DNA damage in BRCA2-deficient cells were further increased by olaparib, which thereby potentiated the innate immune response. Our results also suggest that the impact of PARP inhibitors on the viability of BRCA2-deficient cells may be STAT1-dependent, as STAT1 siRNA-mediated inhibition reduced olaparib toxicity in the short term. Whether STAT1 signaling is also required for the long-term anti-tumoral effect of PARP inhibitors remains to be established.

Our discovery that the chronic response to BRCA2 inactivation is associated with innate immune response upregulation, potentiated by PARP inhibitors, has important therapeutic implications. First, it predicts that drugs that specifically kill BRCA2-deficient cells and tumors by inflicting DNA damage, may concomitantly trigger cytotoxic immune responses. To what extent the deleterious effects of these drugs entail immune response activation remains to be established. Second, it is possible that BRCA-compromised tumors, which become olaparib-resistant through restoration of DNA repair^38–40^ lose the capacity to mount innate immune responses. This tumor subset may be effectively targeted through therapeutic approaches that restore immune signaling.

## METHODS

### Cell lines and growth conditions

Human non-small cell lung carcinoma H1299 cells and human invasive ductal breast cancer MDA-MB-231 cells, wild type (American Type Culture Collection) or carrying a doxycycline (DOX)-inducible BRCA2 shRNA^4^, were cultivated in monolayers in DMEM medium (Sigma) supplemented with 10% tetracycline free foetal bovine serum (Clontech). For induction of shBRCA2, 2 μg/mL DOX (D9891, Sigma) was added to growth medium. Human colorectal adenocarcinoma DLD1 cells, parental and *BRCA2*-mutated (Horizon Discovery^3^), were cultivated in monolayers in DMEM medium (Sigma) supplemented with 10% foetal bovine serum (Thermo Fisher Scientific) and 1% penicillin/streptomycin (Sigma).

### Immunoblotting

To prepare whole cell extracts, cells were washed once in 1xPBS, harvested by trypsinization, washed in 1xPBS and re-suspended in SDS-PAGE loading buffer, supplemented with 0.1 mM DTT. Samples were sonicated and heated at 70°C for 10 min. Equal amounts of protein (50-100 μg) were analysed by gel electrophoresis followed by Western blotting. NuPAGE-Novex 4-12% Bis-Tris and NuPAGE-Novex 3-8% Tris-Acetate gels (Life Technologies) were run according to manufacturer’s instructions.

### RNA-seq analyses

H1299, H1299+shBRCA2^DOX^, MDA-MB-231 and MDA-MB-231+shBRCA2^DOX^ were cultured in presence or absence of DOX for 4 or 28 days. Cells were collected for RNA extraction and processed using the RNeasy^®^ Mini Kit (Qiagen, #74104) according to the manufacturer’s guidelines. RNA samples were quantified using RiboGreen (Invitrogen) on the FLUOstar OPTIMA plate reader (BMG Labtech) and the size profile and integrity analyzed on the 2200 or 4200 TapeStation (Agilent, RNA ScreenTape). RNA integrity number estimates for all samples were between 9.0 and 10.0. Input material was normalized to 1 μg prior to library preparation. Poly(A) transcript enrichment and strand specific library preparation were performed using TruSeq Stranded mRNA kit (Illumina) following manufacturer’s instructions. Libraries were amplified (15 cycles) on a Tetrad (Bio-Rad) using in-house unique dual indexing primers^41^. Individual libraries were normalized using Qubit and size profile was analyzed on the 2200 or 4200

### TapeStation

Individual libraries were pooled together and pooled libraries were diluted to ~10 nM for storage. Each library aliquot was denatured and further diluted prior to loading on the sequencer. Paired end sequencing was performed using a HiSeq4000 75bp platform (Illumina, HiSeq 3000/4000 PE Cluster Kit and 150 cycle SBS Kit), generating a raw read count of >22 million reads per sample.

### RNA-seq data processing

Reads were aligned to the human reference genome (GRCh37) using HISAT2^42^ and duplicate reads removed using the Picard ‘MarkDuplicates’ tool (Broad Institute). Reads mapping uniquely to Ensembl-annotated genes were summarised using featureCounts^43^. The raw gene count matrix was imported into the R/BioConductor environment^44^ for further processing and analysis. Genes with low read counts (less than ~10 reads in more than 3 samples) were filtered out, leaving sets of 14,000 to 15,000 genes to test for differential expression between conditions, depending on the samples considered.

### Differential expression analysis

The analyses were carried out using the R package DESeq2 version 1.18.1^45^. Unless otherwise stated, differentially expressed genes were identified based on two criteria: FDR (False Discovery Rate using Benjamini-Hochberg adjusted *p*-values) < 0.05 and absolute value of Log2(Fold Change) > 0.5. Differentially expressed genes were determined independently for the following human cell lines: DLD1 cells (parental vs *BRCA2*-mutated), H1299, MDA-MB-231, H1299+shBRCA2^DOX^ and MDA-MB-231+shBRCA2^DOX^, +DOX vs −DOX conditions and two time points (4 days and 28 days). For each set of conditions, hierarchical clustering was performed for the top 500 differentially expressed genes using Euclidean distances. Intersections were used to identify differentially expressed genes common to various data sets.

### Gene set enrichment analysis

The GO and REACTOME pathway enrichment analyses were performed using the Gene Set Enrichment Analysis (GSEA) software^46^ with the Molecular Signatures Database collection. The NetworkAnalyst platform^17^ and the Cytoscape 3.5 software^47^ were used to visualize the first-order protein-protein interactions networks, based on the high-confidence (confidence score > 0.9 and experimental evidence required) STRING interactome database^18^.

### Pathway deregulation scores

The Pathifier algorithm^19^ calculates a pathway deregulation score (PDS) based on gene expression matrices for each sample. PDS represents the extent to which the activity of a given pathway differs in a particular test sample from the activity in the matched control. Gene sets were retrieved from the REACTOME database^48^ in GMT format and the R Bioconductor pathifier package version 1.16.0, along with custom R scripts were used to calculate deregulation scores of immune system related pathways in DLD1 *BRCA2^−/−^* cells.

### TCGA data analysis

The Cancer Genome Atlas (TCGA) ovarian serous cystadenocarcinoma annotated mutation files were retrieved from the cBioPortal database (http://www.cbioportal.org/)^49^ for 585 samples respectively. We analyzed mRNA expression by downloading 309 ovarian cases with RNA expression data using the R Bioconductor TCGAbiolinks package version 2.9.5^50^. Samples were split into *BRCA2*-deleted (*BRCA2^del^*) and bulk groups. We determined that there were 4 *BRCA2* deep deletion cases within this cohort. The bulk samples were ranked according to *BRCA2* mRNA levels, allowing us to define a *BRCA2^high^* subgroup (3rd highest quantile of mRNA expression) of 71 samples. Differential expression analysis was performed in the *BRCA2^del^* versus *BRCA2^high^* groups.

### siRNAs

Cells were transfected using Dharmafect 1 (Dharmacon). Briefly, 0.8 x 10^6^ cells were transfected with 40 nM siRNA (ON-TARGET plus human STAT1 SMART pool; L-003543-00-0005, Dharmacon) by reverse transfection in 10-cm plates.

### Cell viability assays

Cells were seeded in 96-well plates at a density previously estimated to reach 70 - 80% confluency after four days in the absence of treatment. Twenty four hours after seeding, drugs were added to the growth medium and viability was determined using resazurin-based assays three days later. Cells were incubated with 10 μg/mL resazurin (R7017, Sigma-Aldrich) diluted in growth medium at 37°C for 2 hours. Fluorescence was measured (544 nm excitation and 590 nm emission) using POLARstar Omega plate reader (BMG Labtech).

### Quantitative PCR (qPCR)

For RNA reverse transcription, the Ambion Kit Power SYBR™ Green Cells-to-CT™ Kit (#4402954) was used according to manufacturer’s instructions. Resulting complementary DNA was analyzed using SYBR Green technology in the QuantStudio 5 Real-Time PCR System. After normalization to *RPL11* or *β-actin*, gene expression was calculated^51^ relative to untreated control cells, as 2^−ΔΔCT^. The following primer pairs were used: *IFIT1*, forward TAC CTG GAC AAG GTG GAG AA and reverse GTG AGG ACA TGT TGG CTA GA; *IFIT2*, forward TGT GCA ACC TAC TGG CCT AT and reverse TTG CCA GTC CAG AGG TGA AT; *IFIT3*, forward CTT CAG TAT TTA CTT GAG GCA GAC and reverse CTT GGT GAC CTC ACT CAT GAC; *IFI6*, forward TCG CTG ATG AGC TGG TCT GC and reverse ATT ACC TAT GAC GAC GCT GC; *OAS1*, forward CGC CTA GTC AAG CAC TGG TA and reverse CAG GAG CTC AGG GCA TA; *OAS2*, forward TCC AGG GAG TGG CCA TAG and reverse TCT GAT CCT GGA ATT GTT TTA AGT C; *ISG15*, forward GCG AAC TCA TCT TTG CCA GTA and reverse CCA GCA TCT TCA CCG TCA G; *TNFRSF1B*, forward CGG GAG CTC AGA TTC TTC CC and reverse GTC TCC AGC TGT GAC CGA AA; *RPL11*, forward GCA AAC TCT GTT CAA CAT CTG and reverse CAT ACT CCC GCA CTT TAG AC; β-actin, forward ATT GGC AAT GAG CGG TTC and reverse GGA TGC CAC AGG ACT CCA T.

### EdU incorporation in asynchronous cells

To label replicated DNA, cells were incubated with 25 μM EdU for 30 min. Samples were collected by trypsinization and fixed using 90% methanol. Incorporated EdU was detected using the Click-iT EdU Alexa Fluor 647 Flow Cytometry Assay Kit (C10634, Invitrogen) according to manufacturer’s instructions. Cells were re-suspended in PBS containing 20 μg/mL propidium iodide (P4864, Sigma) and 400 μg/mL RNase A (12091-021, Invitrogen). Samples were processed using flow cytometry (BD FACSCalibur, BD Biosciences). 10,000 events were analyzed per condition using FlowJo software.

### Antibodies

The following antibodies were used for immunoblotting: mouse monoclonal antibodies raised against BRCA2 (1:1,000, OP95, Calbiochem), GAPDH (1:30,000, 6C5, Novus Biologicals); rabbit polyclonal antibodies raised against STING (1:1,000, CST13647, Cell Signaling), IRF3 (1:1,000, AB76409, Abcam), phospho-IRF3 (1:1,000, AB76493, Abcam), STAT1 (1:1,000, 9175, Cell Signaling), phospho-STAT1 (1:1,000, 9167, Cell Signaling), SMC1 (1:5,000, A300-055A, Bethyl Laboratories).

### *In vivo* xenograft experiments

CB17/SCID mice male (6 weeks old and weighing 26-28 g) were purchased from Charles River Laboratories (Calco, Italy). All animal procedures were in compliance with the national and international directives (D.L. March 4, 2014, no. 26; directive 2010/63/EU of the European Parliament and of the council; Guide for the Care and Use of Laboratory Animals, United States National Research Council, 2011). MDA-MB436 BRCA1-deficient cells, were injected into the mammary fat pad of CB17/SCID female mice. When the tumors were palpable (seven days after cell injection) the treatment was initiated. Each experimental group included five mice. Talazoparib (0.33 mg/kg/day) was administered orally for five consecutive days, followed by two-day break and five more days of treatment. Tumors were excised and processed for RNA extraction using TRIzol reagent (Ambion, Life Technologies).

## Supporting information

## ACKNOWLEDGEMENTS

We thank the Oxford Genomics Centre at the Wellcome Centre for Human Genetics (funded by Wellcome Trust grant reference 203141/Z/16/Z) for the generation and initial processing of the sequencing data. We are grateful to Dr. Andrew Blackford (Department of Oncology, University of Oxford) for critical reading of the manuscript and to Dr. Johanna Michl (DPAG, University of Oxford) for help with RNA-seq analyses of DLD1 cells. Research in M.T. laboratory is supported by Cancer Research UK, Medical Research Council, University of Oxford and EMBO Young Investigator Program. This project has received funding from the European Union’s Horizon 2020 research and innovation programme under the Marie Skłodowska-Curie grant agreement No. 722729.

## AUTHOR CONTRIBUTIONS

M.T., T.R. and E.P.L. designed the RNA-seq study. T.R. performed the RNA-seq experiments, B.W. and H.L. conducted the RNA-seq data alignment, E.P.L. and A.L. conducted analyses of the RNA-seq data. T.R., A.M. and A.H. planned and performed validation of RNA-seq hits, including quantitative RT-PCR for time course analyses. F.G. performed time course analyses of EdU staining and Western blotting. M.P. and A.B. carried out the experiments involving the *in vivo* orthotopic model. M.T. wrote the manuscript.

---

**Supplementary Figure 1.**
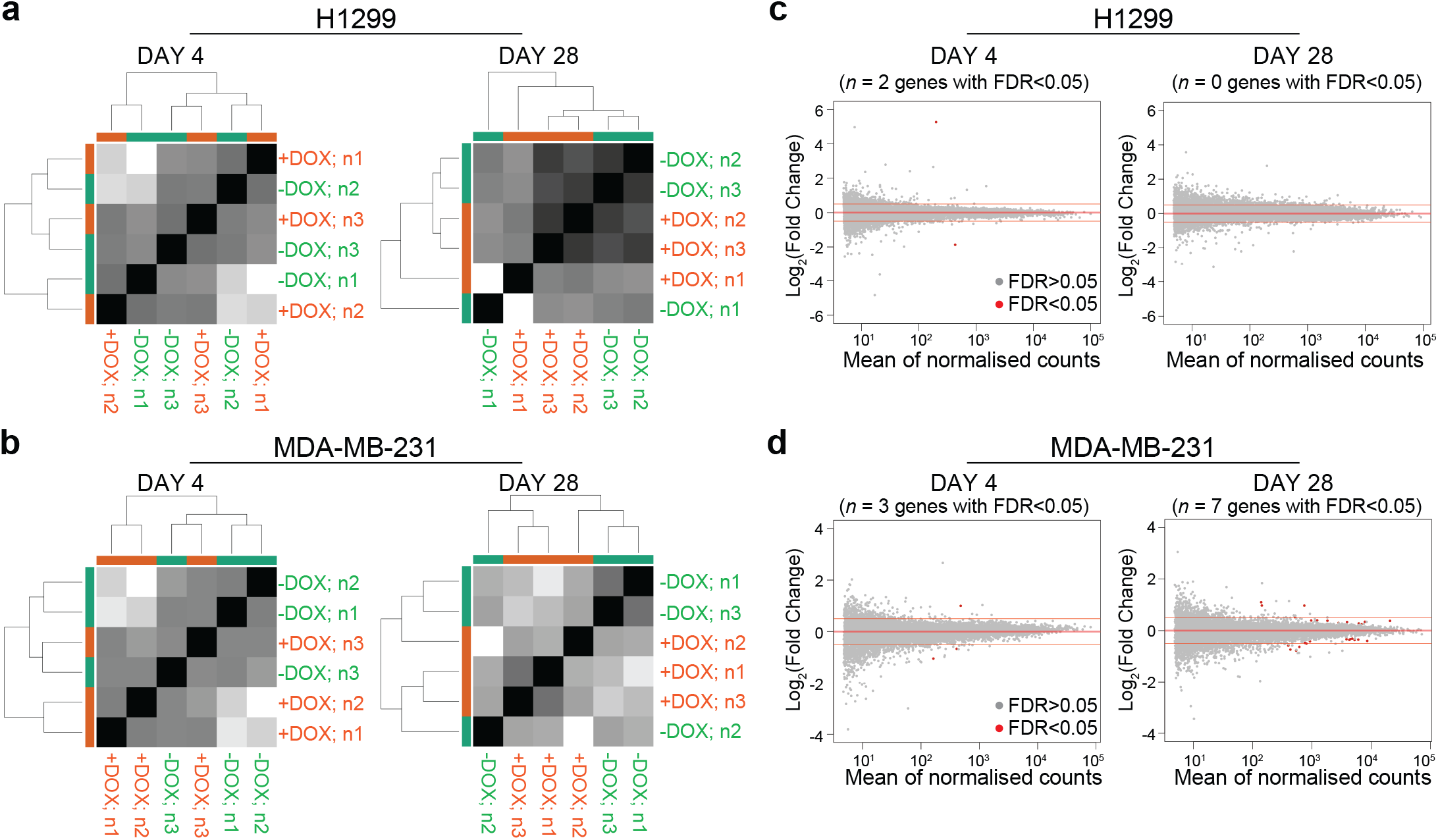
Doxycycline (DOX) treatment does not induce transcriptional changes in parental H1299 or MDA-MB-231 human cells. (**a, b**) Euclidean sample-to-sample distances calculated from RNA-seq normalised count data for H1299 (**a**) and MDA-MB-231 (**b**) cells treated with 2 μg/ml DOX for 4 or 28 days. Hierarchical clustering of distance matrices shows no clear distinction between +DOX and −DOX samples. (**c, d**) MA plots showing differentially expressed genes between +DOX and −DOX samples treated as in (**a, b**). Each dot represents one gene; red, genes with significantly altered expresssion values (FDR < 0.05); gray, genes without significantly altered expresssion values (FDR > 0.05).

**Supplementary Figure 2.**
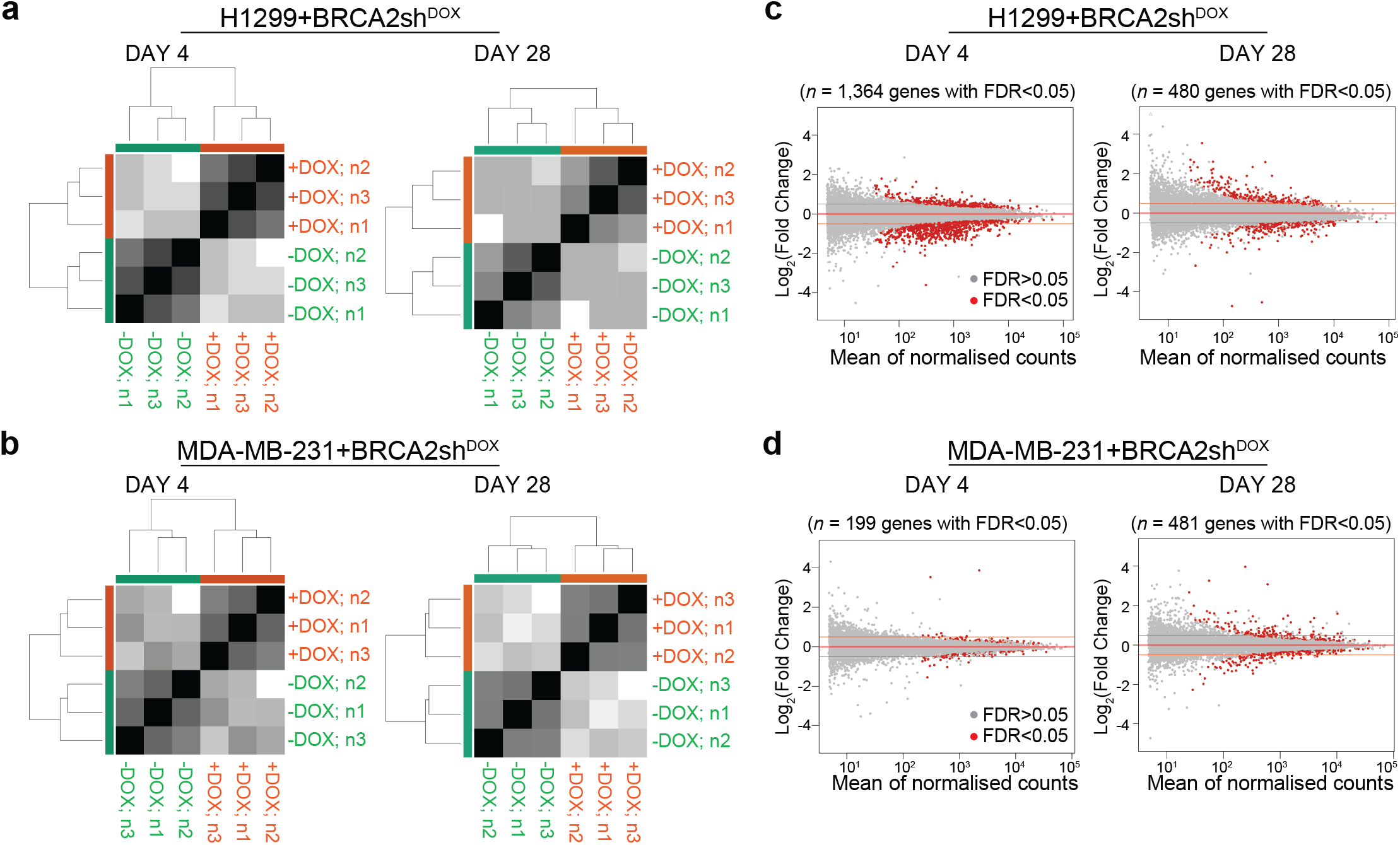
Doxycycline (DOX) treatment induces significant transcriptional changes in H1299 or MDA-MB-231 human cells expressing a DOX-inducible shRNA against *BRCA2*. (**a, b**) Euclidean sample-to-sample distances calculated from RNA-seq normalized count data for H1299 (**a**) and MDA-MB-231 (**b**) cells carrying a DOX-inducible shRNA against BRCA2, treated with 2 μg/ml DOX for 4 or 28 days. Hierarchical clustering of distance matrices shows a clear distinction between −DOX and +DOX samples. (**c, d**) MA plots showing differentially expressed genes between +DOX and −DOX samples treated as in (**a, b**). Each dot represents one gene; red, genes with significantly altered expresssion values (FDR < 0.05); gray, genes without significantly altered expresssion values (FDR > 0.05).

**Supplementary Figure 3.**
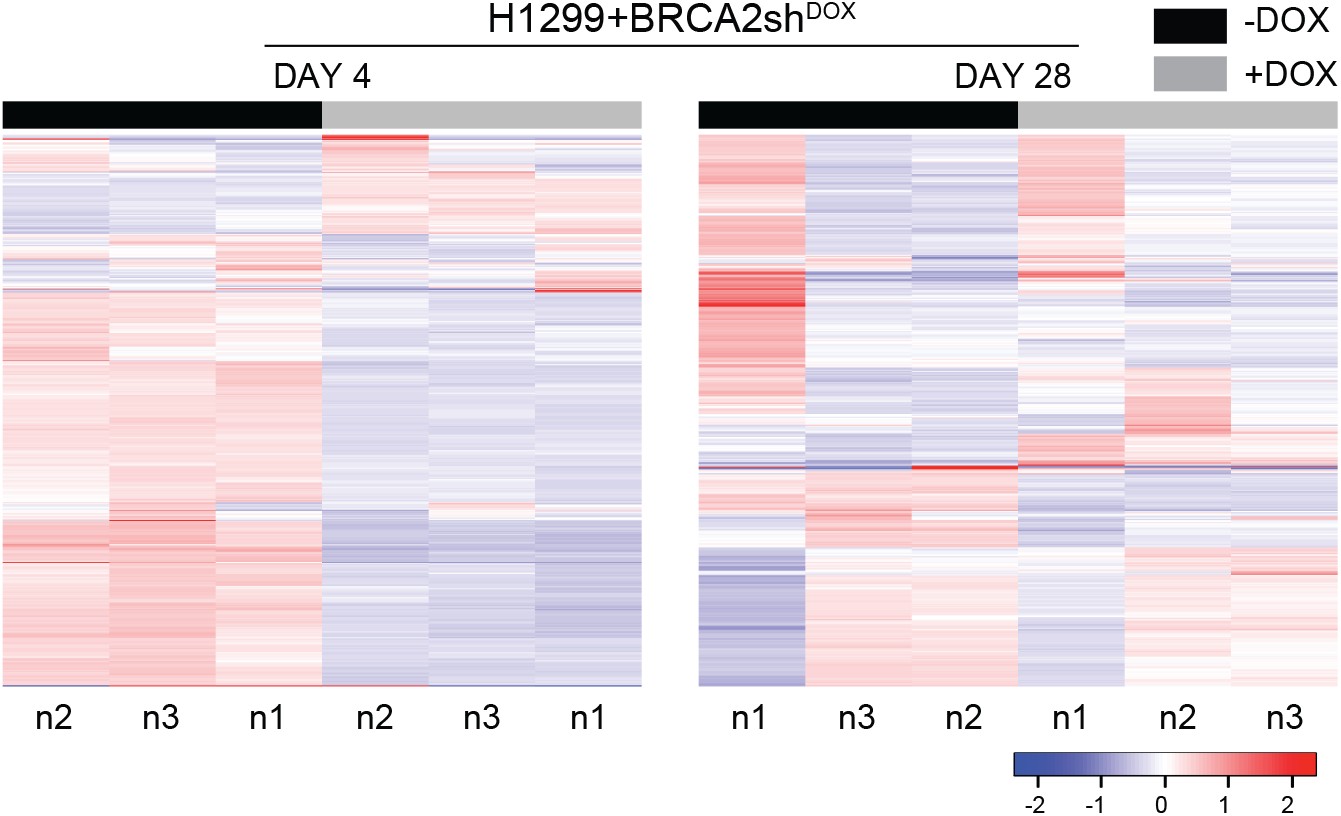
Supervised clustering of log-transformed gene counts obtained from three independent RNA-seq experiments performed in H1299 cells expressing a DOX-inducible shRNA against *BRCA2*. Top scoring 500 genes are shown.

**Supplementary Figure 4.**
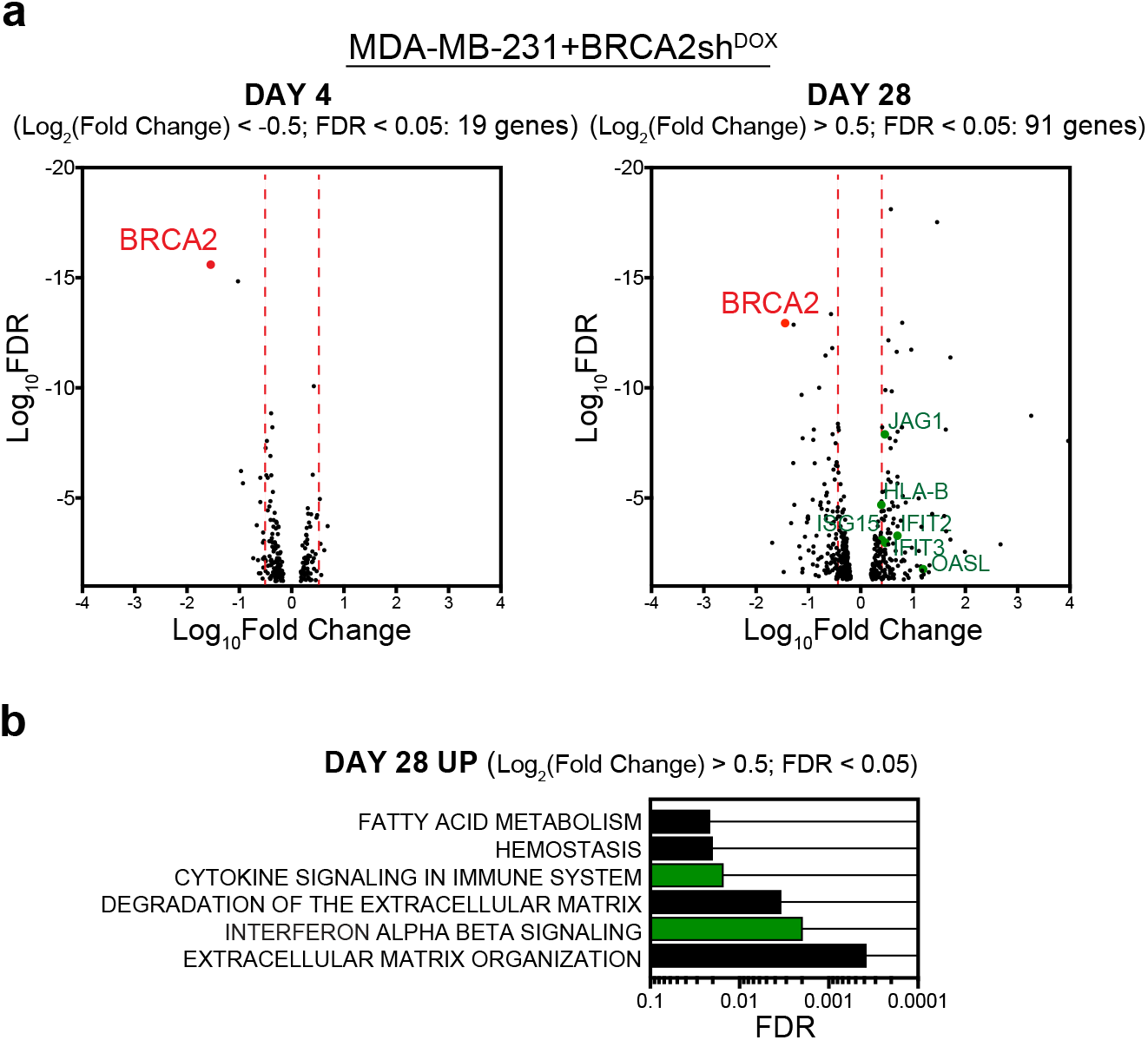
Differential gene expression analyses of MDA-MB-231 cells carrying a DOX-inducible shRNA targeting *BRCA2*. (**a**) Volcano plot of genes differentially expressed (FDR < 0.05) in BRCA2-deficient (+DOX) versus BRCA2-proficient (-DOX) cells, assessed at day 4 or day 28 after DOX addition. (**b**) REACTOME pathway-based enrichment analysis of genes upregulated (Log_2_(Fold Change) > 0.5; FDR < 0.05) in +DOX relative to −DOX samples following 28 days of DOX treatment.

**Supplementary Figure 5.**
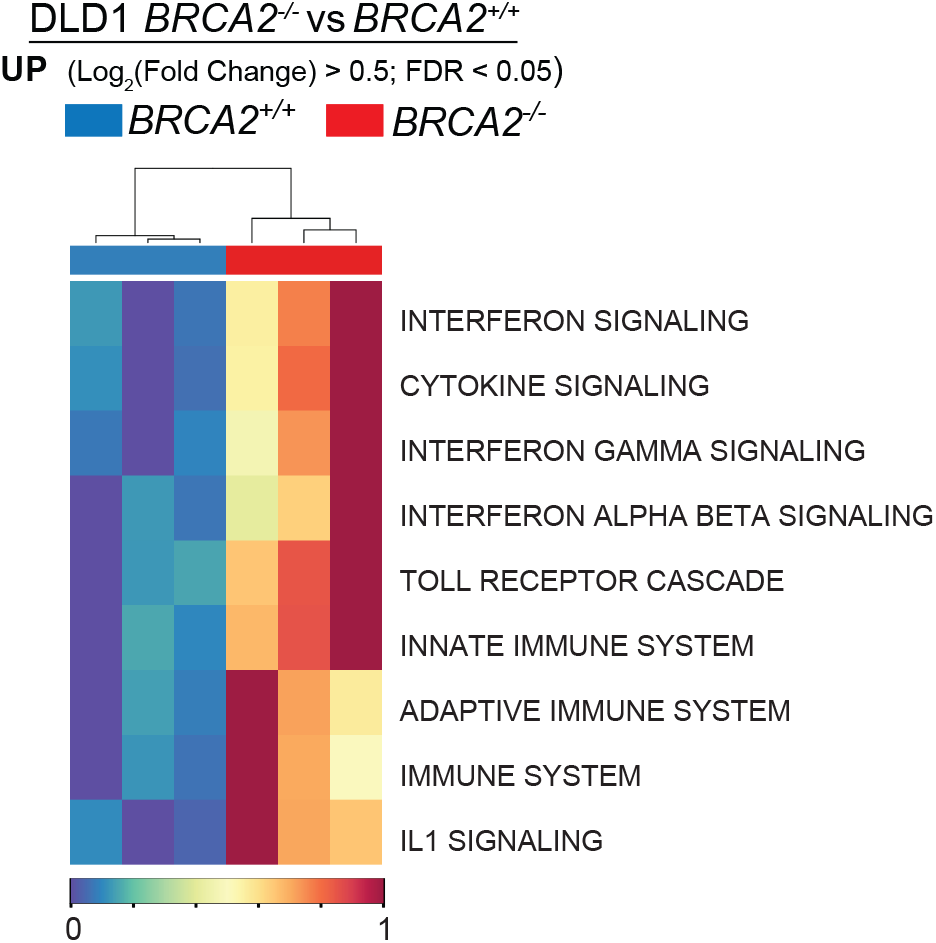
Pathway deregulation score analysis for genes significantly upregulated (Log_2_(Fold Change) > 0.5; FDR < 0.05) in BRCA2^−/−^ vs BRCA2^+/+^ DLD1 cells. Each row corresponds to a Reactome pathway and each column to an RNA-seq sample.

**Supplementary Figure 6.**
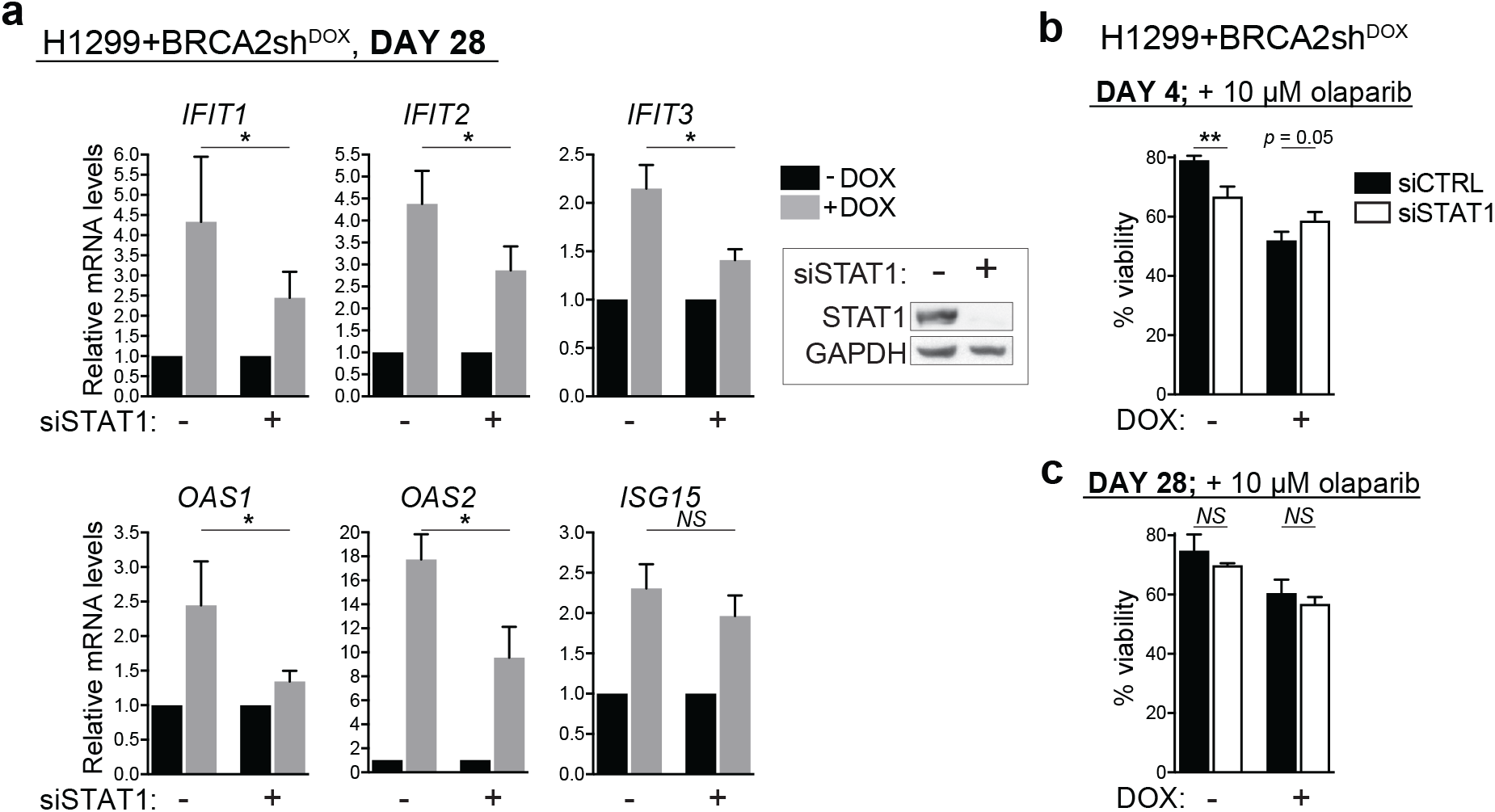
(a) Upregulation of the innate immune response genes triggered by BRCA2 inactivation is STAT1-dependent. H1299 cells expressing a doxycycline (DOX)-inducible BRCA2 shRNA were grown in the presence or absence of DOX for 28 days. Cells transfected with STAT1 siRNA for 48 hours were subjected to quantitative RT-PCR analyses using primers specific for the indicated genes. mRNA levels were expressed relative to the gene encoding β-actin and to untreated (-DOX) controls (2^−ΔΔCT^). Error bars represent SD of *n* = 3 independent experiments, each in performed in triplicate. *, *p* < 0.05 (unpaired two-tailed t test). Whole cell extracts were immunoblotted as indicated. (**b, c**) Effect of STAT1 on viability of BRCA2-deficient cells treated with olaparib. Cells were treated with DOX for 4 days (**b**) or 28 days (**c**) and transfected with indicated siRNAs before treatment with olaparib (10 μM) for 72 hours. Cell viability was determined using resazurin-based assays and reported relative to solvent-treated cells. Error bars represent SD of *n* = 3 independent experiments, each performed in triplicate. *NS, p* > 0.05; *, *p* < 0.05; **, *p* < 0.01 (unpaired two-tailed *t* test).

**Supplementary Figure 7.**
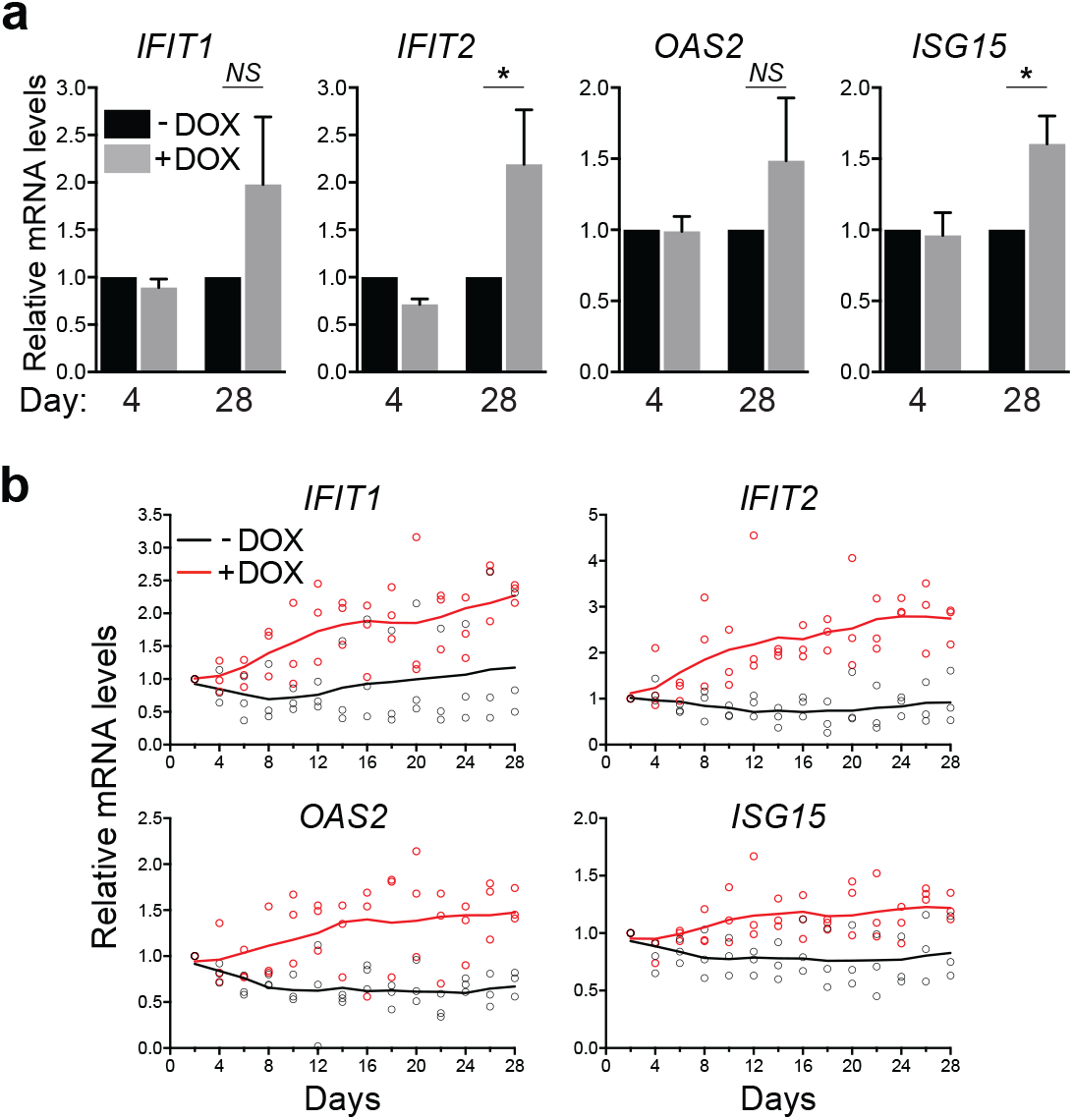
Upregulation of innate immune response genes upon DOX-induced BRCA2 abrogation in MDA-MB-231 cells. (**a**) MDA-MB-231 cells expressing a doxycycline (DOX)-inducible BRCA2 shRNA were grown in the presence of DOX for 4 or 28 days. mRNA levels of indicated genes were determined using quantitative RT-PCR analyses and were expressed relative to the housekeeping gene RPL11 and to untreated (-DOX) control cells. Error bars represent SD of *n* = 3 independent experiments, each in performed in triplicate. *NS, p* > 0.05; *, *p* < 0.05 (paired two-tailed t test). (**b**) MDA-MB-231 cells were cultured as in (**a**) and samples collected every two days. mRNA levels of indicated genes were determined using quantitative RT-PCR analyses and were expressed relative to the gene encoding β-actin and to untreated (-DOX) control cells. Data were obtained from *n* = 3 independent experiments, each in performed in triplicate.

